# Whole-genome analyses resolve the phylogeny of flightless birds (Palaeognathae) in the presence of an empirical anomaly zone

**DOI:** 10.1101/262949

**Authors:** Alison Cloutier, Timothy B. Sackton, Phil Grayson, Michele Clamp, Allan J. Baker, Scott V. Edwards

## Abstract

Palaeognathae represent one of the two basal lineages in modern birds, and comprise the volant (flighted) tinamous and the flightless ratites. Resolving palaeognath phylogenetic relationships has historically proved difficult, and short internal branches separating major palaeognath lineages in previous molecular phylogenies suggest that extensive incomplete lineage sorting (ILS) might have accompanied a rapid ancient divergence. Here, we investigate palaeognath relationships using genome-wide data sets of three types of noncoding nuclear markers, together totalling 20,850 loci and over 41 million base pairs of aligned sequence data. We recover a fully resolved topology placing rheas as the sister to kiwi and emu + cassowary that is congruent across marker types for two species tree methods (MP-EST and ASTRAL-II). This topology is corroborated by patterns of insertions for 4,274 CR1 retroelements identified from multi-species whole genome screening, and is robustly supported by phylogenomic subsampling analyses, with MP-EST demonstrating particularly consistent performance across subsampling replicates as compared to ASTRAL. In contrast, analyses of concatenated data supermatrices recover rheas as the sister to all other non-ostrich palaeognaths, an alternative that lacks retroelement support and shows inconsistent behavior under subsampling approaches. While statistically supporting the species tree topology, conflicting patterns of retroelement insertions also occur and imply high amounts of ILS across short successive internal branches, consistent with observed patterns of gene tree heterogeneity. Coalescent simulations indicate that the majority of observed topological incongruence among gene trees is consistent with coalescent variation rather than arising from gene tree estimation error alone, and estimated branch lengths for short successive internodes in the inferred species tree fall within the theoretical range encompassing the anomaly zone. Distributions of empirical gene trees confirm that the most common gene tree topology for each marker type differs from the species tree, signifying the existence of an empirical anomaly zone in palaeognaths.

## Introduction

The scaling-up of multigene phylogenetic data sets that accompanied rapid advances in DNA sequencing technologies over the past two decades was at first heralded as a possible end to the incongruence resulting from stochastic error associated with single-gene topologies (Rokas et al. 2003, Gee 2003). However, it soon became clear that conflicting, but highly supported, topologies could result from different data sets when sequence from multiple genes was analyzed as a concatenated supermatrix, leading Jeffroy et al. (2006) to comment that phylogenomics - recently coming to signify the application of phylogenetic principles to genome-scale data (Delsuc et al. 2005) - could instead signal ‘the beginning of incongruence’. On the one hand, these observations highlighted the need for more sophisticated models to account for nonphylogenetic signal such as convergent base composition or unequal rates that can become amplified in large concatenated data sets (Jeffroy et al. 2006). But at the same time, there was a growing recognition that gene trees and species trees are not equivalent entities and that horizontal gene transfer, gene duplication and loss, and incomplete lineage sorting can result in heterogeneous topologies for gene trees that are all contained within a single overarching species tree history (Maddison 1997). When gene tree discordance arising from these biological factors predominates, concatenation approaches, which implicitly assume that all genes share the same topology, can not only fail to recover the true species tree but can also infer erroneous relationships with strong support (Kubatko and Degnan 2007, Roch and Steel 2015). This recognition of the biological basis for gene tree discordance prompted proposals for a conceptual shift to adopt the multispecies coalescent model as a flexible and biologically realistic framework that incorporates population-level processes into phylogenetic inference (Degnan and Rosenberg 2009, Edwards 2009, Edwards et al. 2016).

Among the processes generating variation in gene tree histories, coalescent variation caused by incomplete lineage sorting (ILS) has received much attention, in part because it is the most ubiquitous and common form of gene tree-species tree incongruence (Edwards 2009). Incomplete lineage sorting occurs when ancestral polymorphisms do not reach fixation between successive divergences among taxa or, viewed from the opposite direction as deep coalescence, when alleles of two taxa fail to coalesce in their most recent common ancestor (Maddison 1997, Degnan and Rosenberg 2009). The likelihood of ILS increases when internal branches in the species tree are short relative to the effective population size and this phenomenon is therefore likely to accompany many rapid radiations that are the focus of evolutionary studies (Song et al. Liu et al. 2015b). A surprising outcome of ILS under the multispecies coalescent is that there exist branch lengths in the species tree for which the most probable gene tree topology differs from that of the species tree, producing a zone where these anomalous gene trees (AGTs) predominate in the gene tree distribution (the ‘anomaly zone’, Degnan and Rosenberg 2006). The anomaly zone carries clear implications for phylogenetic inference, since any method employing a democratic vote among loci will fail in this region of tree space (Degnan and Rosenberg 2006, 2009; Huang and Knowles 2009). Incomplete lineage sorting is documented for many taxa (e.g. Prüfer et al. 2012, Song et al. 2012, Suh et al. 2015), and the theoretical basis of the anomaly zone is well established (Degnan and Rosenberg 2006, 2009; Rosenberg 2013). However, an empirical anomaly zone has thus far only been reported in Scincidae skinks (Linkem et al. 2016) and the general relevance of this phenomenon to empirical studies has been questioned (Huang and Knowles 2009).

Short branches that produce high amounts of incomplete lineage sorting are also expected to carry relatively few informative substitutions (Huang and Knowles 2009, Xi et al. 2015). Observed gene tree heterogeneity accompanying an inferred empirical anomaly zone could therefore reflect uninformative gene trees rather than truly anomalous ones (Huang and Knowles 2009). More generally, gene tree estimation error has been raised as a potential concern for current methods of species tree inference even outside of the anomaly zone (Liu et al. 2015c, Roch and Warnow 2015). Fully coalescent single-step Bayesian methods that coestimate gene trees with the species tree, despite providing the most seamless statistical environment for testing phylogenetic hypotheses using the coalescent (Xu and Yang 2016), are as yet computationally prohibitive for large data sets (Liu et al. 2015c, Edwards 2016a), motivating the development of many two-step methods that use estimated gene trees as input for species tree inference (reviewed in Liu et al. 2015a, Edwards 2016a). These summary methods can show good performance under simulation (Liu et al. 2010, Mirarab et al. 2016), sometimes even within the anomaly zone (Liu et al. 2010), but assume that gene trees are known without error. Simulation studies have shown that summary species tree methods can still perform well in the presence of gene tree estimation error (e.g. Liu et al. 2010, Mirarab et al. 2016), and recent results from coalescent simulations (Xi et al. 2015) and analysis of empirical data (Blom et al. 2017) indicate that the accuracy of species tree inference can be improved by adding more loci, even if those loci are minimally informative (Xu and Yang 2016). However, whether these observations apply broadly across empirical data sets is as yet unknown, and it is also crucial that gene tree error, if it occurs, is unbiased (Xi et al. 2015). It is therefore imperative for empirical studies to carefully consider the underlying gene tree support for inferred species trees and to assess consistency of results across analysis methods and data types.

Here, we use genome-wide data sets of three types of noncoding markers (conserved non-exonic elements [CNEEs], introns, and ultraconserved elements [UCEs]) to investigate relationships within the Palaeognathae, one of the two basal lineages of modern birds (the other being Neognathae, which includes the Galloanserae [gamebirds + waterfowl] and Neoaves [all other birds]). Palaeognaths encompass the flighted tinamous (Tinamiformes) of South and Central America and the flightless ratites, including the African ostrich (Struthioniformes), Australian emu and Australasian cassowaries (Casuariiformes), New Zealand kiwi (Apterygiformes), and South American rheas (Rheiformes) as well as the recently extinct New Zealand moa (Dinornithiformes) and Madagascan elephant birds (Aepyornithiformes). Palaeognath phylogenetic relationships have remained controversial for over a century (reviewed in Houde 1988, Cracraft et al. 2004), and recent molecular studies have added to the debate by placing the tinamous as sister to the moa within a paraphyletic ratite clade (Phillips et al. 2010, Haddrath and Baker 2012, Baker et al. 2014) and recovering an unexpected sister-group relationship between kiwi and elephant birds (Mitchell et al. 2014, Grealy et al. 2017, Yonezawa et al. 2017). These findings have important implications for our understanding of Gondwonan biogeography and the possibility that morphological convergence has accompanied multiple independent losses of flight in the ratites (Harshman et al. 2008, Haddrath and Baker 2012, Baker et al. 2014, Sackton et al. 2018).

Although ratite paraphyly is strongly supported by all recent molecular studies (Hackett et al. 2008, Harshman et al. 2008, Phillips et al. 2010, Haddrath and Baker 2012, Smith et al. 2013, Baker et al. 2014, Mitchell et al. 2014, Grealy et al. 2017, Yonezawa et al. 2017), rheas have been variously placed as the sister to all other non-ostrich palaeognaths (e.g. Hackett et al. 2008, Harshman et al. 2008, Phillips et al. 2010, Mitchell et al. 2014, Prum et al. 2015 and others), to tinamous (Harshman et al. 2008, Smith et al. 2013; note that moa sequences were absent from these analyses), or to a clade containing emu, cassowary, and kiwi (Haddrath and Baker 2012, Prum et al. 2015), with the same data sometimes producing conflicting results under different analysis regimes (e.g. Harshman et al. 2008, Smith et al. 2013, Prum et al. 2015). Alternative placements of rheas are often accompanied by low bootstrap support, and it is suspected that ILS across short internal branches separating major palaeognath lineages could underlie some of the difficulties in resolving phylogenetic relationships within this group (Haddrath and Baker 2012).

Newly available whole-genome sequences (Zhang et al. 2014, Le Duc et al. 2015, Sackton et al. 2018) allow investigation of palaeognath relationships at a phylogenomic scale. Recently, we reported fully congruent results using the coalescent species tree method MP-EST (Liu et al. 2010) that recovered rheas as the sister to emu/cassowary + kiwi. These results were consistent across marker type and robust to the effects of missing data, alignment uncertainty, and outgroup choice, but differed from the incongruent results obtained under concatenation (Sackton et al. 2018). Here, we corroborate the species tree topology inferred from sequence-based analysis with genome-wide screening for informative presence/absence insertions of CR1 retroelements, and incorporate results from a second species tree method (ASTRAL, Mirarab and Warnow 2015) to further strengthen support for this topology relative to that obtained from concatenated sequence data. Additionally, we employ phylogenomic subsampling to investigate consistency in the underlying signal for conflicting relationships recovered from species tree versus concatenation approaches, and use likelihood evaluation and coalescent simulation to assess the underlying gene tree support for the recovered species tree topology and the existence of an empirical anomaly zone in palaeognaths. Throughout, we consider the variation in signal among classes of noncoding nuclear markers that are becoming increasingly adopted for genome-scale analyses (Edwards et al. 2017), and contrast the relative performance of these markers under different analysis regimes to resolve historically challenging relationships among Palaeognathae.

## Materials & Methods

### Data set compilation

We assembled data sets for three types of noncoding nuclear markers: conserved non-exonic elements (CNEEs), introns, and ultraconserved elements (UCEs) for 14 palaeognath species and a chicken outgroup from publicly available whole-genome sequence assemblies (Suppl. Table S1). We chose to analyze noncoding sequences primarily because coding regions across large taxonomic scales in birds are known to experience more severe among-lineage variation in base composition than noncoding regions, which can complicate phylogenetic analysis (Jarvis et al. 2014, Reddy et al. 2017). On the other hand, noncoding markers can be more challenging to align than coding markers, and the three marker types we analyze here exhibit a range of variation and alignment challenges (Edwards et al. 2017).

We filtered the 1,949,832 conserved elements (CEs) identified statistically by Sackton et al. (2018) to omit elements overlapping annotated exon, gene, or CDS features in the galGal4 chicken genome release (NCBI annotation release 102) to generate a candidate set of 811,696 conserved non-exonic elements (CNEEs). This set was further filtered to retain 16,852 elements greater than 250 bp in length. Orthologous sequence from 10 newly published palaeognath genomes (Sackton et al. 2018) as well as ostrich *(Struthio camelus)* and white-throated tinamou *(Tinamus guttatus)* genomes from the Avian Phylogenomics Project (Zhang et al. 2014) was compiled by lifting over reference genome coordinates from chicken to each target palaeognath in an existing 42 species multiway whole-genome alignment (WGA, Sackton et al. 2018) with HAL Tools v.2.1 (Hickey et al. 2013). Liftover output was parsed to retain only uniquely mapping regions between the chicken reference and target palaeognath genomes. CNEEs with no missing taxa from the 12 palaeognaths included in the whole-genome alignment, and with at least as many variable sites as taxa were retained (N= 14,528 loci). Loci overlapping the set of UCEs described below were omitted using BEDTools v. 2.26.0 (Quinlan and Hall 2010), leaving a data set of 12,676 CNEEs.

Candidate introns were identified using BEDTools to output coordinates for annotated intron features in the galGal4 genome release that did not overlap with any annotated exon feature. Chicken coordinates for these introns were lifted over to target palaeognaths in the whole-genome alignment as described for CNEEs above, filtered to remove duplicated regions, and expected exon/intron boundaries in each palaeognath sequence were refined by making intron coordinates flush to liftovers for adjacent exon sequences. Candidate introns within each palaeognath species were omitted if shorter than 100 bp, if longer than 100 kb, or if longer than 10 kb and more than 50% longer than the chicken reference intron length. After adding sequence from the published North Island brown kiwi (Le Duc et al. 2015) and little bush moa (Cloutier et al. 2018) as described below, one intron per gene was chosen, requiring a minimum pairwise sequence identity of 70% and fewer than 0.5 undetermined sites (e.g. gaps and Ns) per bp aligned sequence, and then choosing among multiple introns per gene based on the smallest number of missing taxa and longest average unaligned input sequence length across taxa. This produced a final data set of 5,016 introns, each originating from a different gene.

Ultraconserved elements (UCEs) were compiled using the set of 3,679 bioinformatically harvested UCEs from the Avian Phylogenomics Project (Jarvis et al. 2014, 2015; accessed from http://dx.doi.org/10.5524/101041). In addition to chicken, we used ostrich and white-throated tinamou included in the Avian Phylogenomics Project data as reference species to lift over to all other palaeognaths in the whole-genome alignment. Liftover output from these multiple reference taxa was parsed to omit duplicated regions and tiled for regions that were consistent across reference species. A maximum of one missing taxon per locus from the 12 palaeognaths in the whole-genome alignment was allowed, resulting in a data set of 3,158 UCEs.

Blastn searches with NCBI’s default ‘somewhat similar’ parameters (evalue 1e-10, perc_identity 10, penalty -3, reward 2, gapopen 5, gapextend 2, word size 11) were used with query sequence from each of the three kiwi species included in the whole-genome alignment to identify orthologous regions in the North Island brown kiwi *(Apteryx mantelli*, Le Duc et al., which was not included in the WGA. North Island brown kiwi sequence was added for loci that had consistent top hits across blastn searches, with a single high-scoring segment pair (HSP) covering at least 50% of the input query sequence and minimum 80% sequence identity. Using this approach, we added *A. mantelli* sequence for 11,534/12,676 CNEEs (91% of total), 2,560/5,016 introns (51%), and 2,033/3,158 UCEs (64%).

Sequence was also added from a reference-based genome assembly of the extinct little bush moa *(Anomalopteryx didiformis*, Cloutier et al. 2018) that was generated by mapping moa reads to the emu genome included in the whole-genome alignment. Emu coordinates from the liftover approach outlined above were used to retrieve the corresponding region in moa, and moa sequence was retained if it spanned at least 30% of the emu reference length or was at least 200 bp in length, excluding Ns. Moa sequence was added for 12,643/12,676 CNEEs (99.7%), 4,982/5,016 introns (99.3%), and 3,146/3,158 UCEs (99.6%).

Sequences for each locus were retrieved from genome assemblies and aligned *de novo* with MAFFT v. 7.245 (Katoh and Standley 2013) using default options for global iterative alignment for CNEEs (option ‘ginsi’) and local iterative alignment for introns and UCEs (option ‘linsi’). This produced data sets of 4,794,620 bp of aligned sequence for the 12,676 CNEEs, 27,890,802 bp for 5,016 introns, and 8,498,759 bp for 3,158 UCEs.

### Gene tree inference

Best maximum likelihood trees were built with RAxML v. 8.1.5 (Stamatakis 2014) from unpartitioned alignments for each locus using a GTR+GAMMA substitution model and 20 independent searches beginning with randomly generated starting trees. Topologies were also inferred for 500 bootstrap replicates per locus, with a GTR+GAMMA model on unpartitioned data in RAxML.

### Species tree inference

MP-EST v. 1.5 (Liu et al. 2010) analyses were conducted with three full runs per data set beginning with different random number seeds and with ten independent tree searches within each run. The species tree topology was inferred using best maximum likelihood gene trees as input, and node support was estimated from MP-EST runs using gene tree bootstrap replicates as input.

ASTRAL-II v. 4.10.9 (Mirarab and Warnow 2015, hereafter ‘ASTRAL’) was also run using the best maximum likelihood gene trees to infer the species tree topology, and the 500 gene tree bootstrap replicates to estimate node support. To permit direct comparison across methods, the default quadripartition support values output by ASTRAL were replaced with traditional bipartition supports by plotting output bootstrap replicates from ASTRAL on the inferred species tree topology using RAxML.

To investigate the effects of concatenation on phylogenetic inference, ExaML v. 3.0.16 (Kozlov et al. 2015) was used to infer maximum likelihood topologies from fully partitioned (i.e. 1 partition per locus) concatenated data sets using a GTR+GAMMA substitution model. Twenty-one full tree searches were run for each data set, with 20 searches beginning from complete random starting trees, and one additional search using a random stepwise addition order parsimony starting tree. A minimum of 50 bootstrap replicates were computed for each data set, with additional replicates performed as necessary until convergence was reached as assessed by bootstopping analysis with a majority-rule consensus tree criterion implemented in RAxML.

For all three methods, bootstrap supports were placed on the inferred species tree using RAxML, and trees were outgroup rooted with chicken using ETE v. 3 (Huerta-Cepas et al. 2016).

### Phylogenomic subsampling

In addition to the traditional bootstrapping approaches outlined above, phylogenomic subsampling (Blom et al. 2017; reviewed in Edwards 2016b) was used to assess the consistency of the underlying support for conflicting clades. For each marker type (CNEEs, introns, and UCEs), loci were randomly sampled with replacement to create subsets of 50, 100, 200, 300, 400, 500, 1000, 1500, 2000, 2500, and 3000 loci. This process was repeated ten times to create a total of 110 data sets per marker type (e.g. 10 replicates of 50 loci each, 10 replicates of 100 loci, etc. for each of CNEEs, introns, and UCEs). Topologies were inferred for bootstrap replicates of each data set using MP-EST, ASTRAL, and ExaML as described above. However, for reasons of computational tractability, 200 rather than 500 bootstrap replicates were used for each method (including ExaML), and MP-EST was run once (rather than three times) for each data set, although still with ten tree searches within each run.

Support for alternative hypotheses regarding the sister group to the rheas, and the sister to emu + cassowary, was estimated by first counting the number of bootstraps that recovered each alternative topology from the 200 bootstraps run for each replicate, and then calculating the mean value for each hypothesis across the ten replicates within each data set size category.

### Analysis of incomplete lineage sorting (ILS) and gene tree heterogeneity

Analyses of gene tree heterogeneity were conducted using both the best maximum likelihood trees inferred by RAxML, as well as majority-rule extended consensus trees generated from RAxML bootstrap replicates. Most analyses of topological incongruence required an identical taxon sampling across gene trees; however, there was a much larger amount of missing data for North Island brown kiwi than for other taxa, in part because this species was added *post hoc* to data sets that were compiled for species included in the whole-genome alignment and in part because the assembly quality of this species was lower than for those palaeognaths in the WGA. We therefore pruned North Island brown kiwi from all gene trees, and omitted loci that were missing any other taxa. This approach retained 12,643/12,676 CNEEs (99.7% of loci), 4,982/5,016 introns (99.3%), and 2,866/3,158 UCEs (90.8%).

Gene support frequency (GSF) and internode certainty ‘all’ (ICA, Salichos et al. 2014) were calculated with RAxML. Distributions of gene tree topologies were calculated by computing all pairwise Robinson-Foulds (RF) cluster distances on outgroup rooted gene trees with TreeCmp v. 1.1-b308 (Bogdanowicz et al. 2012) and parsing the output into mutually compatible sets of trees with identical topology where RF= 0. Numbers of substitutions occurring on braches conflicting with the species tree topology were inferred under a parsimony criterion in PAUP v. 4.0a151 (Swofford 2002).

Relative support for each gene tree topology was assessed by computing ΔAIC from the log-likelihood score (lnL) of the inferred gene tree and lnL when the input alignment for each locus was constrained to the species tree topology. Both likelihood values were calculated in RAxML on unpartitioned alignments with a GTR+GAMMA substitution model, and branch lengths were optimized for each topology, a criterion for applying the ΔAIC approach.

Approximately unbiased (AU) tests were run in IQ-TREE v. 1.5.4 (Nguyen et al. 2015). For each locus, we tested the estimated gene tree topology against an *a priori* candidate set of probable trees that enforced monophyly of the five higher-level palaeognath lineages (kiwi, emu + cassowary, rheas, moa + tinamous, ostrich), but allowed all possible rearrangements among those lineages (for a total of 105 trees in the candidate set, plus the gene tree topology itself if it did not occur within this set). We also tested gene trees against a second set of candidate topologies using the same criteria as above, but additionally allowing all possible rearrangements within a monophyletic tinamou clade (for 1,575 candidate topologies). For each gene, in addition to the reported P-value for the fit of the species tree topology, we also calculated the rank of the species tree topology relative to all tested candidates from P-values sorted in ascending order.

The proportion of observed gene tree heterogeneity consistent with coalescent variation was estimated through simulation. For each marker type, we estimated ultrametric species tree branch lengths in mutational units (μΤ, where μ is the mutation rate per generation and T is the number of generations) by constraining concatenated alignments of all loci to the species tree topology with a GTR+GAMMA substitution model and strict molecular clock in PAUP. For this purpose, we ignore the additional mutations incurred due to variation in ancestral lineages in the species tree (Angelis and dos Reis 2015), as well as misestimation of the number of mutations due to deviation of the gene trees from the species tree (Mendes and Hahn 2016). These mutational branch lengths were used in combination with the MP-EST and ASTRAL species tree branch lengths in coalescent units (τ= T/4N_e_) to calculate the population size parameter theta (Θ) for each internal branch following Degnan and Rosenberg (2009), where (Θ/2) · T/N_e_ = μΤ which reduces to Θ= μT/τ. Theta for terminal branches was set to a constant value of 1; this value had no impact on simulated gene trees because we simulated only one allele per species.

10,000 gene trees were simulated from species trees using these mutational branch lengths and theta values for each data set (e.g. CNEEs, introns, and UCEs, with theta derived from both MP-EST and ASTRAL branch lengths) using the sim.coaltree.sp function in Phybase v. 1.5 (Liu and Yu 2010). Pairwise Robinson-Foulds cluster and matching cluster distances of each simulated gene tree to the species tree topology were calculated with TreeCmp (Bogdanowicz et al. 2012). The ratio of mean gene tree-species tree distances for simulated gene trees relative to mean distances for empirically estimated gene trees was calculated as a measure of the amount of observed gene tree heterogeneity that can be accounted for by coalescent processes.

Rooted triplets were generated by pruning all possible combinations of three ingroup taxa plus the chicken outgroup from each gene tree with ETE. Proportions of the major (species tree) triplet and the two alternative minor triplets were calculated for all species trios, as well as for combinations of higher level lineages (e.g. all trios involving one kiwi, one rhea, and either emu or cassowary). We followed the approach of Song et al. (2012) to account for non-independence of sampled trios when calculating proportions for higher-level lineages by assigning the most commonly occurring triplet combination for each gene and omitting genes with ties before tallying triplet counts across all genes. Observed triplet values were compared to expected proportions calculated using branch lengths in coalescent units from MP-EST and ASTRAL species trees, where the expected frequency of the major triplet equals 1 - 2/3 exp(-t) and the two minor triplets equal 1/3 exp(-t), with t representing the internal branch length in the triplet (Pamilo and Nei 1988).

Equation 4 from Degnan and Rosenberg (2006) was used to calculate the value *a(x)* for each internal branch *x* measured in coalescent units from MP-EST and ASTRAL species trees. Calculated *a(x)* values were compared to coalescent branch lengths for each descendent internal branch *y*, with *y* < *a(x)* providing evidence that this region of the species tree falls within the zone where anomalous gene trees (AGTs) are expected - the anomaly zone. In addition to calculations from species tree branch lengths, which are derived from input best maximum likelihood gene trees, we also performed anomaly zone calculations for each of the 500 species tree bootstrap replicates using scripts provided by Linkem et al. 2016 (accessed from http://datadryad.org/resource/doi:10.5061/dryad.sf6s9).

### CR1 retroelement insertions

Patterns of CR1 retroelement insertions were used to corroborate findings from sequence-based analyses for all palaeognath species included in the whole-genome alignment (therefore excluding the North Island brown kiwi and little bush moa). Draft genomes for each species were repeat masked using RepeatMasker v. 4.0.5 (Smit et al. 2015) with RepBase Update 20140131, the sensitive search option, and query species ‘vertebrata metazoa’. RepeatMasker output was parsed to retain simple CR1 insertions at least 50 bp in length, omitting overlapping and nested insertions. Coordinates for up to 500 bp of flanking sequence to either side of CR1 insertions in each species were retrieved and lifted over to every other target palaeognath species in the whole-genome alignment with HAL Tools.

Liftover coordinates were parsed to exclude duplicated regions, and orthologous loci were compiled across species with BEDTools and custom Perl scripts. Loci with sequence from at least one taxon for each major lineage (kiwi, emu + cassowary, rheas, ostrich, and tinamous) and with phylogenetically informative insertion patterns were retained (i.e. excluding symplesiomorphies shared by all palaeognaths or autapomorphic insertions unique to a single species). Sequence for each locus was extracted from genome assemblies and *de novo* aligned with MAFFT using default parameters for global alignment. Presence/absence scores based on RepeatMasker annotations were bioinformatically reassessed in aligned sequence data, and sequences with gaps spanning the CR1 insertion site were rescored as ‘omitted’. Loci with flush insertion boundaries and identifiable target site duplications (TSDs) at least 2 bp in length were retained, and all retained alignments were visually inspected. RepeatMasker output for these loci was parsed to verify that putative orthologous insertions shared the same CR1 subtype, insertion orientation, and degree of fragmentation. Sequence from chicken was added to alignments for CR1 insertions that grouped all non-ostrich palaeognaths to verify insertion boundaries and confirm that CR1 absence was not due to a secondary deletion within this genomic region in ostrich. Statistical support for conflicting patterns of CR1 insertions was calculated with the KKSC insertion significance tests of Kuritzin et al. (2016).

## Results

### Coalescent species tree methods, but not concatenation, recover congruent palaeognath relationships

Using genome-wide data sets of 12,676 CNEEs, 5,016 introns, and 3,158 UCEs, we recover fully congruent topologies across all marker types and for the combined total evidence tree using MP-EST and ASTRAL coalescent species tree approaches (Fig. 1, Suppl. Fig. S1a-h; N.B. following Liu et al. 2015b and Edwards 2016a, we note the distinction between fully coalescent MP-EST and ASTRAL, whose clustering algorithm is not strictly coalescent, but hereafter refer to both methods as ‘coalescent’). Maximal support is obtained throughout for MP-EST and for the ASTRAL species tree built from CNEEs, but with reduced support for the placement of rheas as the sister to a clade containing kiwi and emu + cassowary for ASTRAL trees inferred from introns and UCEs (95% and 83% bootstrap support, respectively). In contrast, ExaML inference from concatenated data sets places the rheas as sister to all other non-ostrich palaeognaths with full support for introns and reduced support for CNEEs (60%) and UCEs (90%, Suppl. Fig. S1i-l). ExaML analyses are also inconsistent across marker types in their placement of a casuariiform clade comprising emu and cassowary, where introns recover a sister group relationship to kiwi as is seen for MP-EST and ASTRAL, but with casuariiforms instead placed as the sister to moa + tinamous with 60% support for CNEEs and 96% support for UCEs (Suppl. Fig. S1).

**Figure 1.**
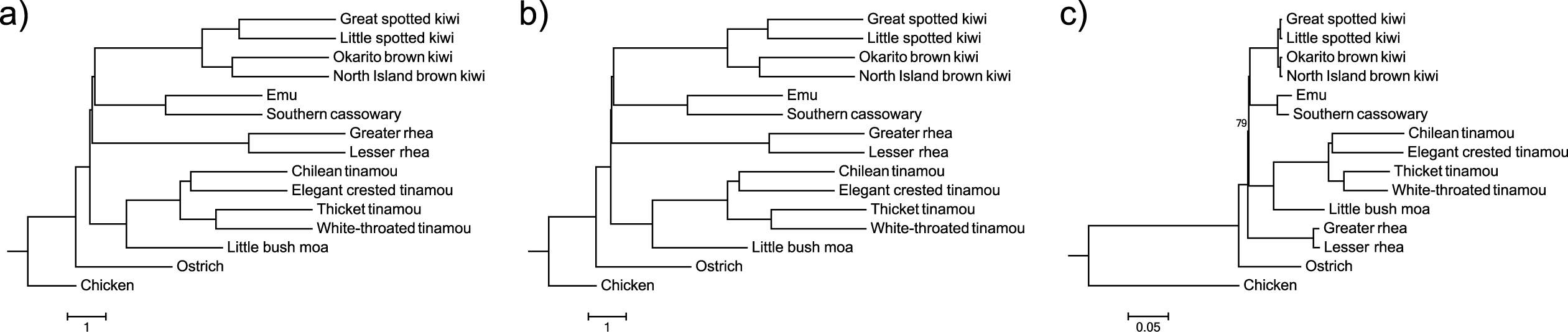
Palaeognath relationships inferred from the total evidence data set (TENT) of 20,850 loci using MP-EST (a) and ASTRAL (b) species tree methods, and maximum likelihood analysis of concatenated data with ExaML (c). Bootstrap support values are indicated for clades with < 100% support. Scale bars indicate branch length in coalescent units for MP-EST and ASTRAL and in substitutions/site for ExaML. Terminal branches in MP-EST and ASTRAL species trees are uninformative and are drawn as a constant value across taxa.

Robustness of the underlying signal for these inconsistently recovered relationships was further assessed with phylogenomic subsampling, which uses a variation of double bootstrapping (Seo et al. 2008) to generate replicate data sets with increasing numbers of randomly sampled loci, each of which is then subject to traditional bootstrap resampling across sites. MP-EST analyses rapidly accumulate support for a sister-group relationship between rheas and emu/cassowary + kiwi, where 100% support is consistently recovered across replicates with an intermediate number of loci (e.g. with > 500 loci, Fig. 2a-c, Fig. 3a,j), and with support for alternative hypotheses sharply dropping off for replicates with greater than 200 loci. Support accumulates more slowly for ASTRAL, but the hypothesis of rheas as sister to emu/cassowary + kiwi clearly dominates and support for alternatives declines in replicates with more than 1000 loci for all markers (Fig. 2d-f, Fig. 3). In contrast, subsampling replicates are less consistent for relationships inferred under concatenation with ExaML (Fig. 2g-i, Fig. 3). In particular, CNEEs oscillate between recovering rheas as the sister to moa + tinamous, or the sister to all other non-ostrich palaeognaths, although always with low bootstrap support (Fig. 2g, Fig. 3g). The other two marker types more clearly support the topology recovered from full data sets with ExaML that place rheas as sister to the remaining non-ostrich palaeognaths, although bootstrap support for UCE replicates is generally weak (Fig. 2h,i; Fig. 3).

**Figure 2.**
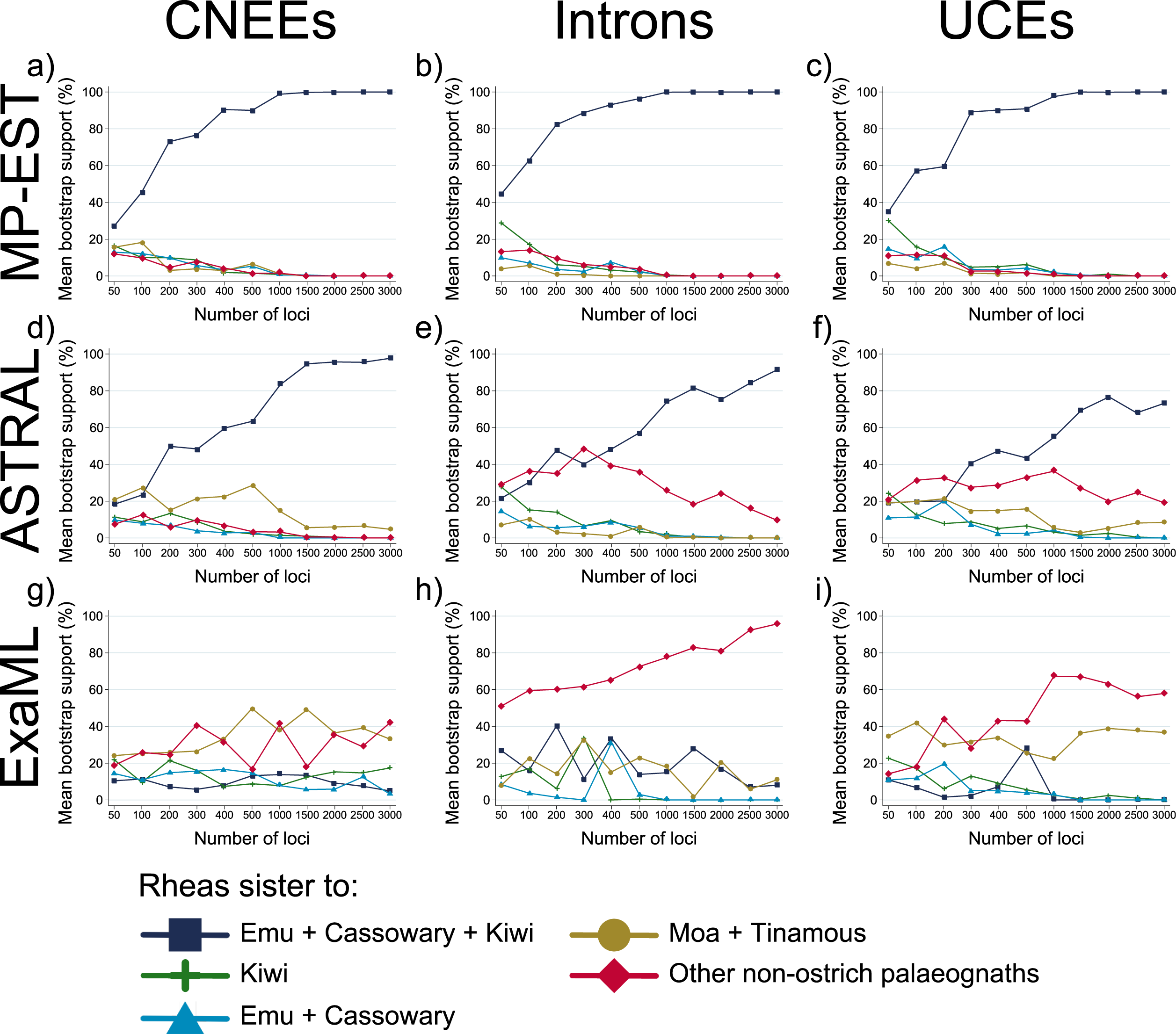
Support for alternative hypotheses of the sister group to rheas from phylogenomic subsampling using MP-EST (a-c), ASTRAL (d-f), and ExaML (g-i). Plots display the mean bootstrap support for each hypothesis from 10 replicates of randomly sampled loci within each data set size category (e.g. 50-3000 loci, shown on x-axis).

**Figure 3.**
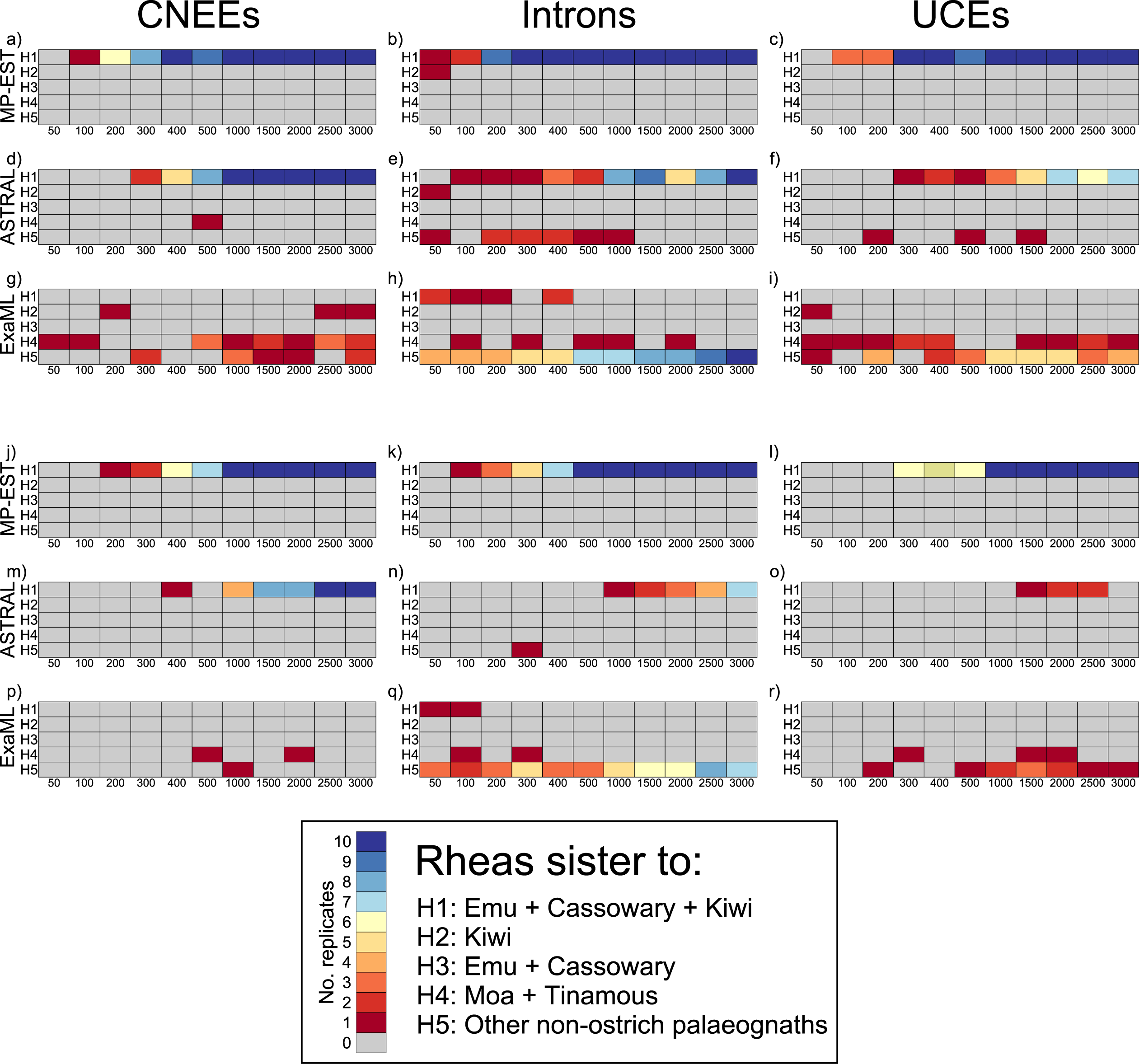
Heatmap of the number of replicates from phylogenomic subsampling that support each alternative hypothesis for the sister group to rheas at a minimum bootstrap support of 70% (a-i) and 90% (j-r). Rows labelled H1-H5 within each panel correspond to the five alternative hypotheses outlined in the legend. Columns within each panel labelled 50-3000 indicate the number of loci subsampled at random with replacement from all loci for each marker type. Coloring of cells indicates the number of replicates (of 10 in total for each data set size category) that support each hypothesis at the given bootstrap cut off, corresponding to the colouring scheme outlined in the legend.

Subsampling provides even more robust support for emu + cassowary as the sister to kiwi, with both MP-EST and ASTRAL quickly accumulating support for this clade and with rapidly declining support for all other hypotheses (Suppl. Fig. S2a-f, Suppl. Fig. S3). ExaML intron replicates also steadily accumulate support for this relationship with an increasing number of loci (Suppl. Fig. S2h, Suppl. Fig. S3h,q). The alternative hypothesis of emu + cassowary as sister to moa + tinamous, which is favored by CNEEs and UCEs analyzed within a concatenation framework, is not well supported by subsampling, where conflicting topologies characterized by low support are recovered across ExaML replicates (Suppl. Fig. S2g,i; Suppl. Fig. S3).

### CR1 retroelement insertions corroborate findings from sequence-based analyses

Patterns of CR1 retroelement insertions corroborate both the inferred species tree topology from MP-EST and ASTRAL and the existence of substantial conflicting signal consistent with incomplete lineage sorting across short internal branches. In total, 4,301 informative CR1 insertions were identified from multispecies genome-wide screens, the vast majority of which (4,274 of 4,301, or 99.4%) are entirely consistent with relationships inferred from sequence-based analyses (Fig. 4a; analysis here was restricted to species in the whole-genome alignment, and little bush moa and North Island brown kiwi are therefore not included). Not surprisingly, we identify many more insertion events occurring along shallower branches with longer estimated lengths and fewer insertions along shorter branches that form the backbone of the inferred species tree (refer to Fig. 1 for estimated branch lengths).

**Figure 4.**
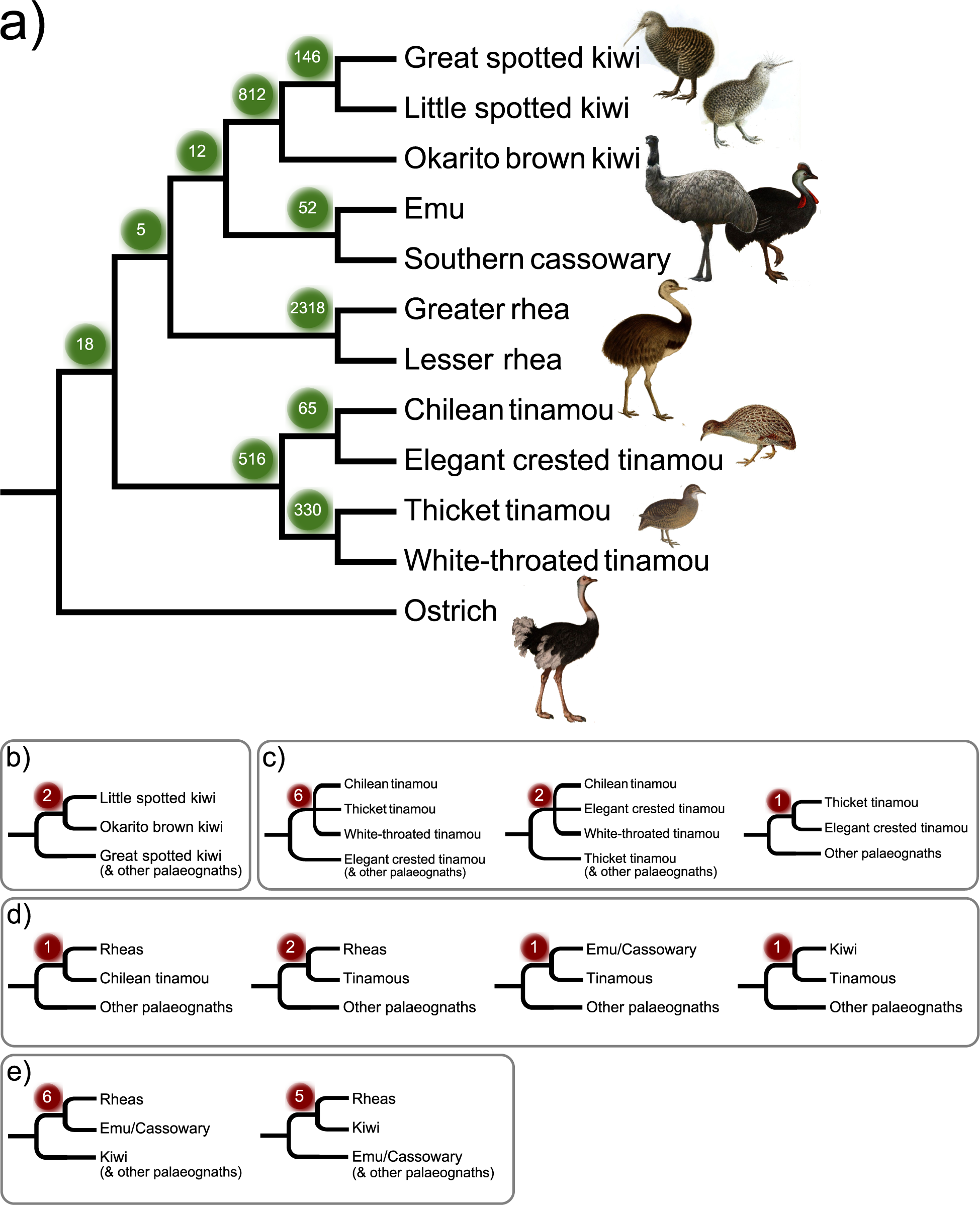
CR1 retroelements corroborate the inferred species tree topology from MP-EST and ASTRAL (a), and also display conflicting insertion patterns consistent with incomplete lineage sorting (b-e). Numbers in colored balls indicate the number of shared retroelement insertions for each clade. Bird illustrations were obtained under the Creative Commons License from Wikimedia Commons (https://commons.wikimedia.org; full details are given in Supplemental Table S2).

Of the 27 (0.6%) CR1s that are inconsistent with the species tree topology, two conflict with the inferred relationships within kiwi, and nine contradict relationships among tinamous (Fig. 4b,c). However, in each case, conflicting CR1s are far outweighed by markers that support the species tree topology, providing robust support from statistical tests of insertion significance (i.e. the KKSC tests of Kuritzin et al. 2016, P= 1.1e^−66^ in support of inferred kiwi relationships, P= 8.8e^−24^ for Chilean tinamou + elegant crested tinamou, and P= 8.4e^−152^ for thicket + white-throated tinamous). A further five insertions are inconsistent with tinamous as sister to a clade containing rheas, kiwi, and emu + cassowary and would instead place tinamous as sister to just one of these other lineages (Fig. 4d). Again, species tree relationships are statistically supported for comparisons of all trios despite these contradictory markers. For example, while casuariiforms (emu and cassowary) and kiwi each share one CR1 insertion exclusively with tinamous, they have 12 insertions shared with each other to the exclusion of tinamous, therefore supporting emu/cassowary + kiwi with P= 8.2e^−05^. Similar calculations support rheas with emu/cassowary to the exclusion of tinamous at P= 0.0018, and rheas with kiwi to the exclusion of tinamous at P= 0.004.

Perhaps most strikingly, we identified six CR1s shared by rheas and emu/cassowary to the exclusion of all other palaeognaths, including kiwi, and five CR1s shared by rheas and kiwi alone (Fig. 4e). Together with the 12 insertions shared by emu, cassowary and kiwi, these patterns are still sufficient to support the inferred relationship of rheas as the sister to an emu/cassowary + kiwi clade (P= 0.048), but indicate substantial conflict in this region of the tree, mirroring the results from sequence-based analysis. Symmetric numbers of CR1s supporting the two conflicting topologies further suggest that ILS, rather than ancestral hybridization, underlies the observed conflicts. Insertion significance tests accordingly support the bifurcating species tree topology rather than a reticulate hybridization network or unresolved polytomy among these three lineages (Kuritzin et al. 2016). We also note that no CR1 insertions were recovered to support the alternative placement of rheas as the sister to all other non-ostrich palaeognaths as produced by concatenation-based analyses (i.e. no CR1s were shared by emu, cassowary, kiwi, and tinamous, but absent from rheas and ostrich).

### Patterns of gene tree heterogeneity suggest substantial incomplete lineage sorting (ILS) and an empirical anomaly zone

We investigated gene tree heterogeneity using both the maximum likelihood estimates of gene trees and the majority rule extended consensus of bootstrap replicates. These analyses produced similar results and are therefore reported for consensus gene trees only.

Distributions of estimated gene tree topologies illustrate that the most common topology for each marker type is not the species tree topology inferred by MP-EST and ASTRAL, thereby suggesting the existence of an empirical anomaly zone (Fig. 5a-c). While the ranking of specific gene tree topologies differs across marker types, common to these anomalous gene trees that occur at higher frequency than the species tree topology is the fact that both the shallowest clades as well as the deepest split between the ostrich and all other palaeognaths are maintained throughout (Fig. 5d). Rearrangements of AGTs relative to the MP-EST and ASTRAL species tree topology instead involve the two short internal branches forming the common ancestor of emu and cassowary with kiwi, and with this clade to rheas (Fig. 5d).

**Figure 5.**
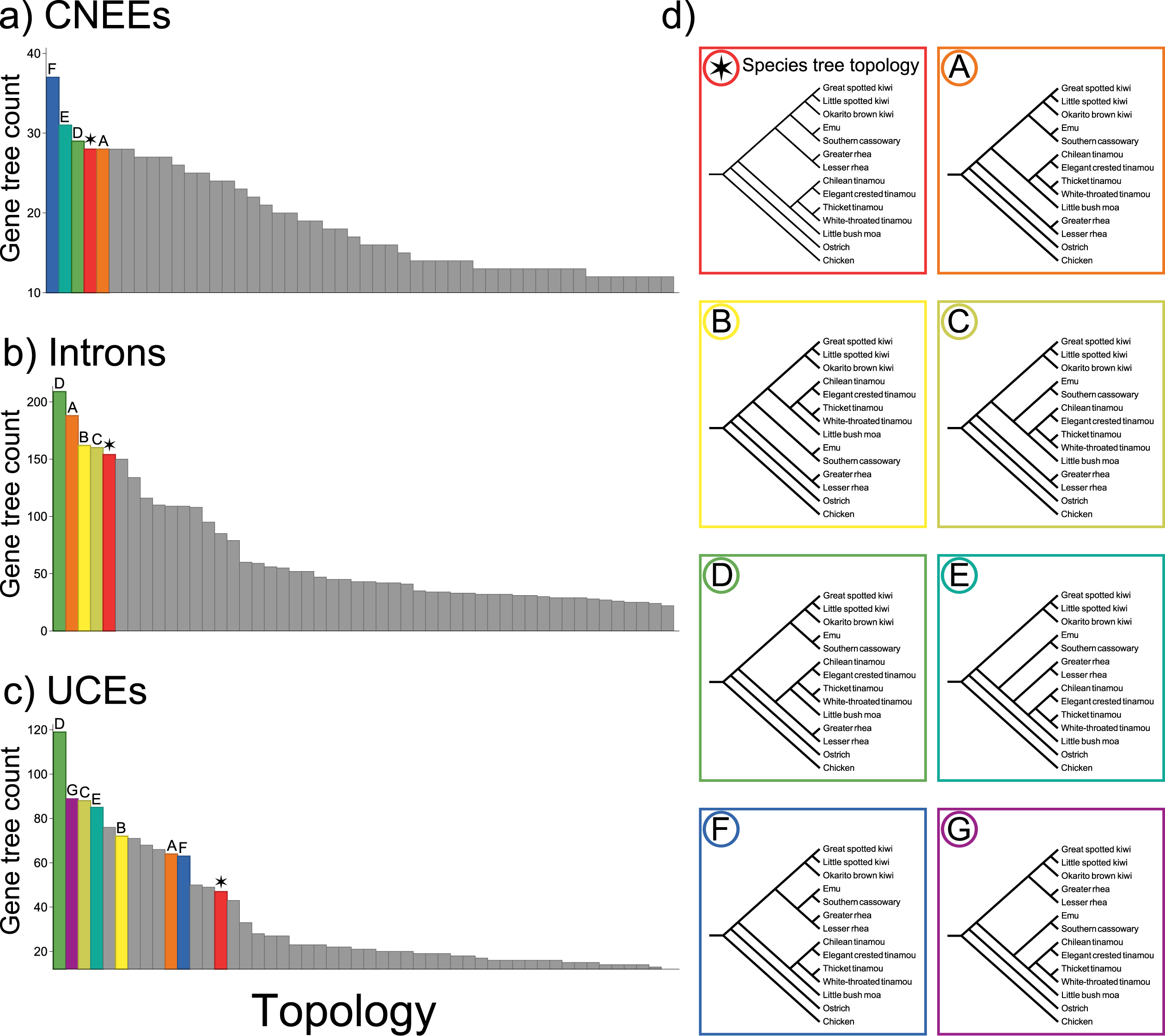
Distributions of the 50 most common gene tree topologies for CNEEs (a), introns (b), and UCEs (c), showing that the most common gene tree topology for each marker type does not match the inferred species tree topology. Symbols and bar coloring in parts a-c correspond to the topologies shown in (d).

To more fully investigate the observed gene tree heterogeneity, we considered all estimated gene trees on a clade-by-clade basis by calculating the gene support frequency (GSF), which tallies the number of genes that recover each bipartition in the inferred species tree, and the internode certainty ‘all’ statistic (ICA, Salichos et al. 2014), which incorporates the frequencies of all commonly occurring bipartitions (Fig. 6). Clades that are recovered consistently in both coalescent and concatenation approaches (e.g. ostrich sister to other palaeognaths, emu + cassowary, moa + tinamous, interrelationships within kiwi, and within tinamous) are also typically supported by the majority of individual gene trees, although singlegene support is weaker for CNEEs (Fig. 6a). However, low (near zero) internode certainty ‘all’ values for emu/cassowary + kiwi indicate that alternative bifurcations occur at roughly equal frequency in the input set of gene trees, and the negative ICA for the relationship of this clade with rheas means that the emu/cassowary + kiwi + rhea clade is actually recovered less often than alternatives.

**Figure 6.**
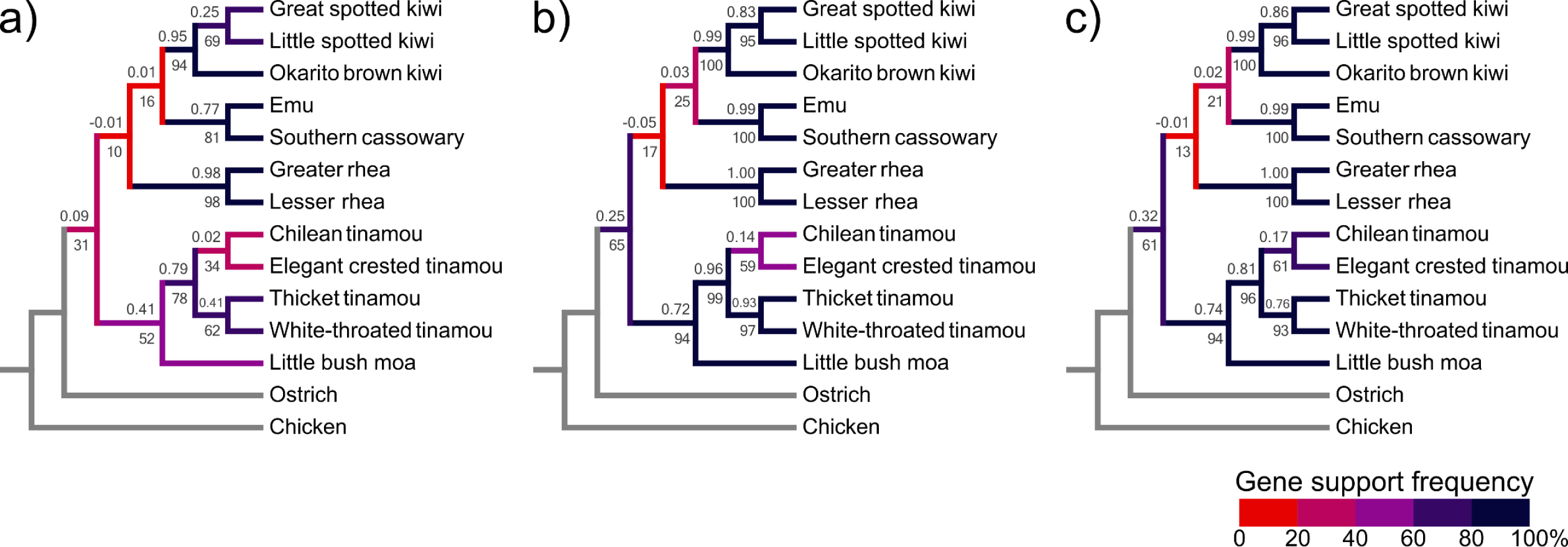
Gene tree support for clades recovered in the MP-EST and ASTRAL species trees for CNEEs (a), introns (b), and UCEs (c). Gene support frequency (GSF, the percent of gene trees containing each clade) is given beneath branches, with heatmap coloring of branches according to GSF as indicated in the legend. Values above branches give the internode certainty ‘all’ statistic (ICA, Salichos et al. 2014), indicating support for each clade relative to all other conflicting bipartitions in the set of gene trees.

We next considered whether topological differences between estimated gene trees and the species tree are well supported, or are instead likely to primarily reflect gene tree estimation error. Mean bootstrap support for estimated gene trees is relatively high, especially for introns and UCEs (83.9% and 82.8% respectively, Fig. 7a-c). However, average support falls by about 10% for each marker type when gene tree bootstrap replicates are constrained to the species tree topology, with P < 0.0001 for paired t-tests of each data set. These results suggest that differences from the species tree are broadly supported by variation in sequence alignments for individual loci. To test this further, we compared the difference in Akaike information criterion (AIC) for estimated gene trees to the AIC obtained when the sequence alignment for each gene was constrained to the species tree topology. Approximately 80% of CNEEs have ΔAIC (gene tree-species tree) less than -2, indicating substantial support in favor of the gene tree topology relative to that of the species tree (Burnham and Anderson 2002), while the proportion was even greater for introns and UCEs (approximately 90% with ΔAIC < -2, Fig. 7d). Despite this result, approximately unbiased (AU) tests typically failed to reject a hypothesis that the data fit the species tree topology, with only about 20% of introns and UCEs, and 30% of CNEEs rejecting the species tree topology at P < 0.05 (Fig. 7d). However, the species tree topology is also not commonly amongst the top 5% of candidate alternative topologies when these alternatives are ranked according to increasing AU test P-value within each locus (Fig. 7d).

**Figure 7.**
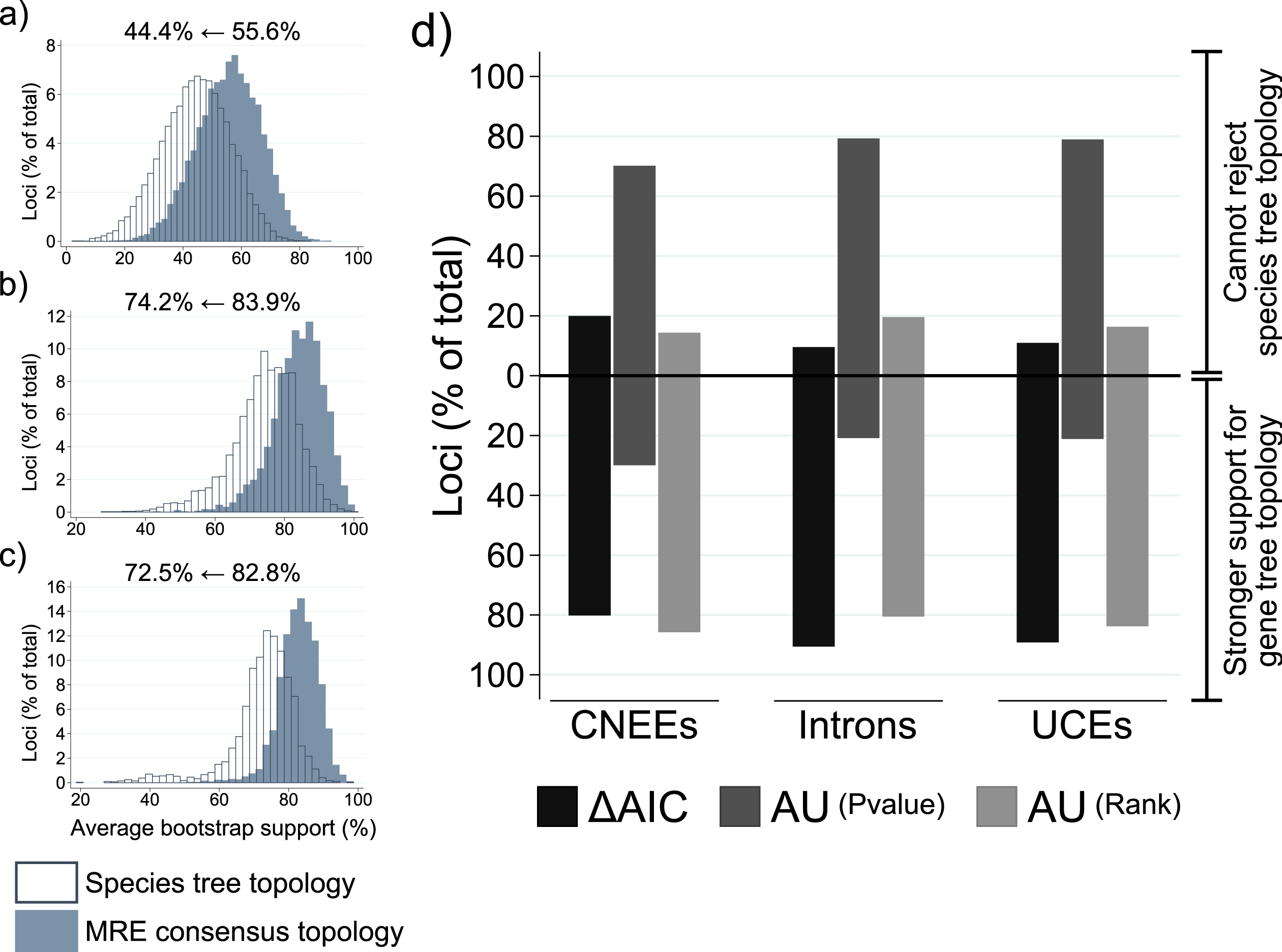
Support for observed topological heterogeneity in estimated gene trees. (a-c) Distributions for the average bootstrap support for all clades recovered in majority rule extended consensus gene trees (‘MRE consensus topology’, drawn as solid bars), and the average support when bootstrap replicates for each gene are constrained to the inferred species tree topology (open bars). Values above each panel indicate the decrease in the grand mean of average support across genes when bootstrap replicates are constrained to the species tree topology. (d) Support that observed gene tree topologies differ from the inferred species tree. Dark gray bars indicate the difference in Akaike information criterion (AIC) from values calculated using the MRE consensus topology for each gene, relative to that obtained when the sequence alignment is constrained to the species tree topology. The baseline at 0 corresponds to ΔAIC = -2, thus values beneath this line indicate loci where likelihood values support the gene tree topology substantially better than that of the species tree. Results of approximately unbiased (AU) tests are indicated with medium gray bars showing the proportion of loci that reject (below baseline) and fail to reject (above baseline) the species tree topology at a P-value cut off of 0.05, and light gray bars showing the proportion of loci where the species tree topology occurs among the top 5% of candidate topologies (above baseline), or is within the bottom 95% of tested topologies (below baseline) for AU P-values ranked in ascending order. AU tests for the 105 and 1575 candidate tree sets produced similar results and are shown for the 105 candidate set only.

In keeping with the results for all loci, gene tree topologies are also generally supported for loci falling within AGT groups. Support for individual gene trees is somewhat weak for CNEEs, with low median bootstrap support and few substitutions occurring along branches that conflict with the species tree topology (Suppl. Fig. S4a,d), which is consistent with the shorter average alignment length and lower variability of these loci (Suppl. Fig. S5). However, support is much stronger for introns and UCEs, with most loci having bootstrap support above 50% for conflicting clades and ΔAIC < -2 indicating much stronger likelihood support for the recovered gene tree topology relative to that obtained if sequence alignments are constrained to match the inferred species tree (Suppl. Fig. S4).

Simulations were used to further assess what proportion of total gene tree heterogeneity is likely attributable to coalescent processes rather than to gene tree estimation error (Fig. 8). Using either Robinson-Foulds (RF) distances or the matching cluster distance, which is influenced less by the displacement of a single taxon than is the RF metric (Bogdanowicz et al. 2012), and simulating from coalescent branch lengths estimated with either MP-EST or ASTRAL for each marker type, we find that coalescent processes alone can account for more than 70% of the observed gene tree heterogeneity in most comparisons, and >90% for introns when gene trees are simulated from MP-EST coalescent branch lengths (Fig. 8).

**Figure 8.**
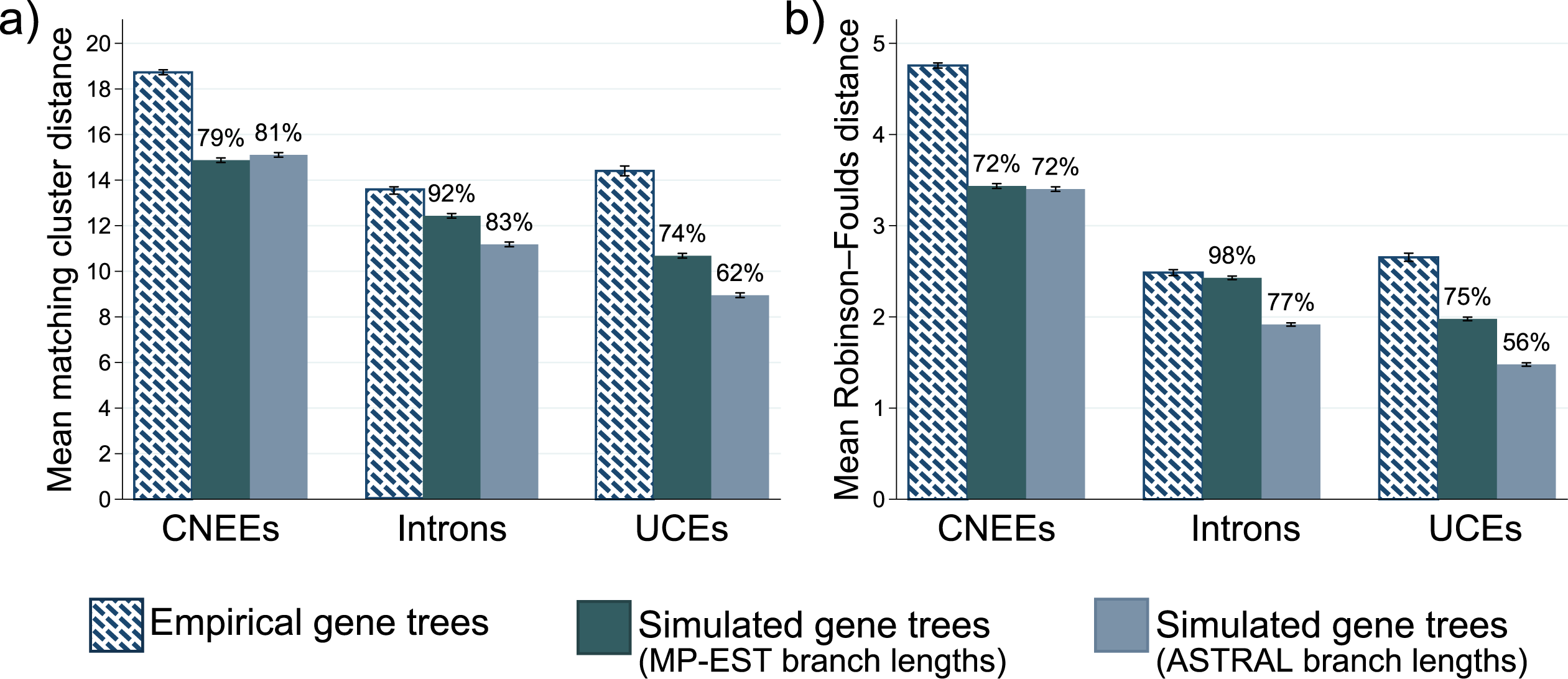
Pairwise distances between gene trees and the MP-EST/ASTRAL species tree topology. Distances between each gene tree and the species tree topology were calculated as the matching cluster distance (a) or Robinson-Foulds cluster distance (b) on outgroup rooted trees, and mean values for all pairwise gene tree-species tree distances within each category are shown. Error bars indicate the 95% confidence interval of the mean. Distances were calculated for empirically estimated gene trees and for data sets of 10,000 gene trees that were simulated using coalescent branch lengths from the MP-EST and ASTRAL species trees. Values above bars for simulated data sets indicate the ratios of means for simulated data sets compared to the mean for empirically estimated gene trees.

### Observed gene tree heterogeneity and anomalous gene trees (AGTs) are consistent with expectations from coalescent theory

Analysis of rooted triplets, where gene trees are decomposed into all possible combinations of three species plus the outgroup, corroborate relationships inferred by MP-EST, which also relies on rooted triplets (Liu et al. 2010) and ASTRAL, which instead uses unrooted quartets (Mirarab and Warnow 2015), with the major, or most frequent, topology for all species trios matching that of the species tree. Observed counts of the major and two minor topologies for the rhea, kiwi, and emu+cassowary lineages (i.e. from sampling all trios that contain one representative of each lineage) also occur in proportions that are consistent with coalescent theory, based on the estimated length of the internal branch within the trio (Pamilo and Nei 1988, Fig. 9 Fisher’s exact P > 0.9 for all comparisons of observed proportions to those expected from either MP-EST or ASTRAL branch lengths). Similar to the results from conflicting CR1 insertions, the two minor topologies for rooted triplets occur at almost equal frequencies, which is consistent with coalescent variation arising from incomplete lineage sorting rather than alternative processes such as introgression. Also notable are the similar triplet proportions observed for all three marker types, which suggest that gene tree estimation error and/or systematic biases within datasets alone are unlikely to explain the observed results.

**Figure 9.**
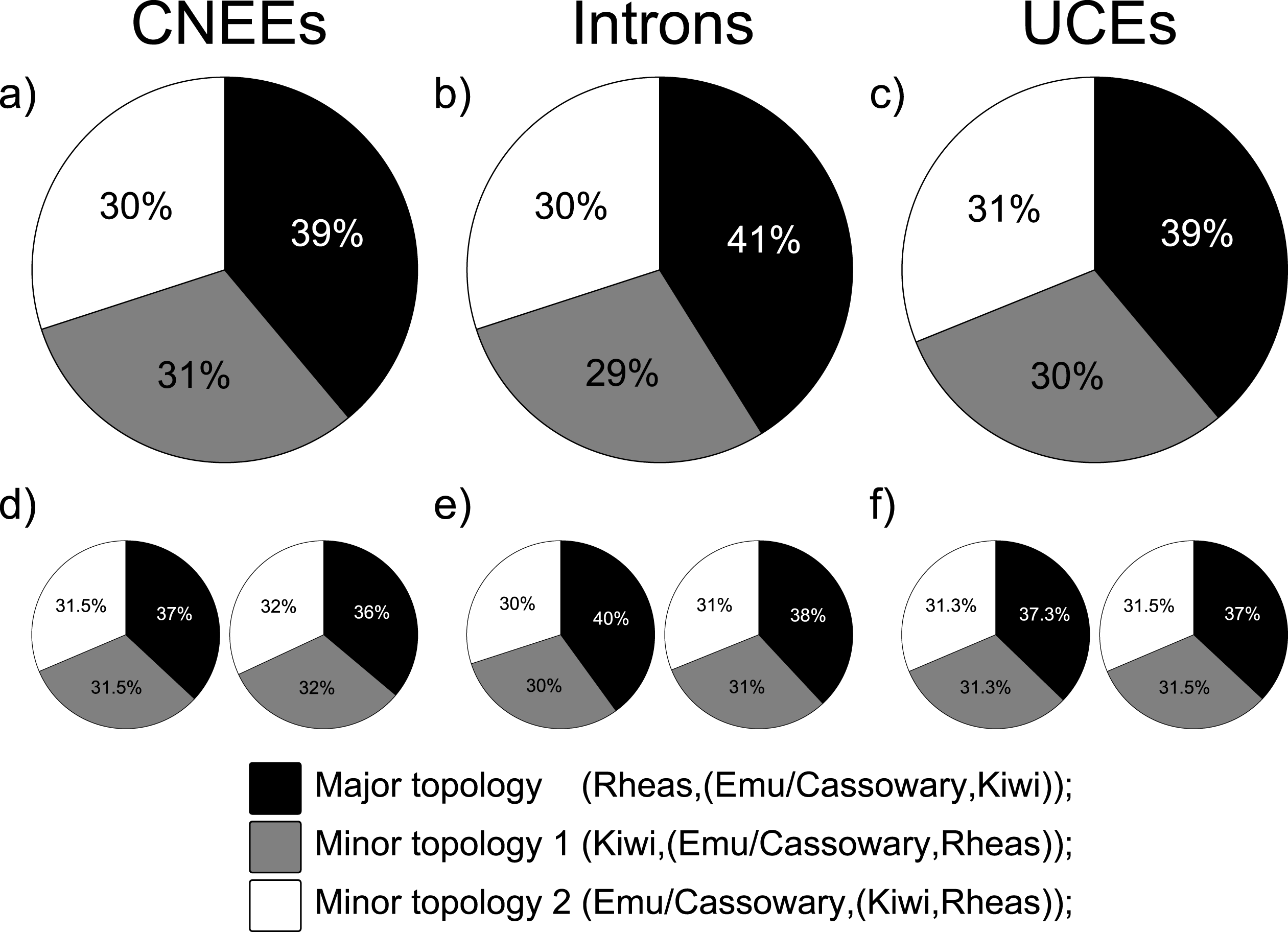
Proportions of rooted triplets for the three alternative topologies involving rheas, kiwi, and emu + cassowary. (a-c) Observed proportions of the major and two minor topologies from empirically estimated gene trees. (d-f) Expected proportions based on the length of the internal branch within the triplet (e.g. the common ancestor of kiwi + emu/cassowary) in coalescent units, following Pamilo and Nei (1988). For d-f, pie charts on the left and right give expected proportions based on branch lengths from MP-EST and ASTRAL species trees, respectively.

The identity and frequency of observed anomalous gene trees are also consistent with expectations from coalescent theory. Following Degnan and Rosenberg (2006) and Rosenberg (2013), branch lengths in coalescent units estimated with either MP-EST (Fig. 10) or ASTRAL (Suppl. Fig. S6) have values expected to produce AGTs across the short successive internal branches separating rheas, emu+cassowary, and kiwi for all marker types. Branch lengths for CNEEs, but not introns or UCEs, are also consistent with AGTs arising across branches separating moa + tinamous from the remaining non-ostrich palaeognaths. Inference of these anomaly zones applied equally when using species tree branch lengths, or for branch lengths from each bootstrap replicate (i.e. every bootstrap replicate that recovers the species tree topology also places these regions of tree space within the anomaly zone).

**Figure 10.**
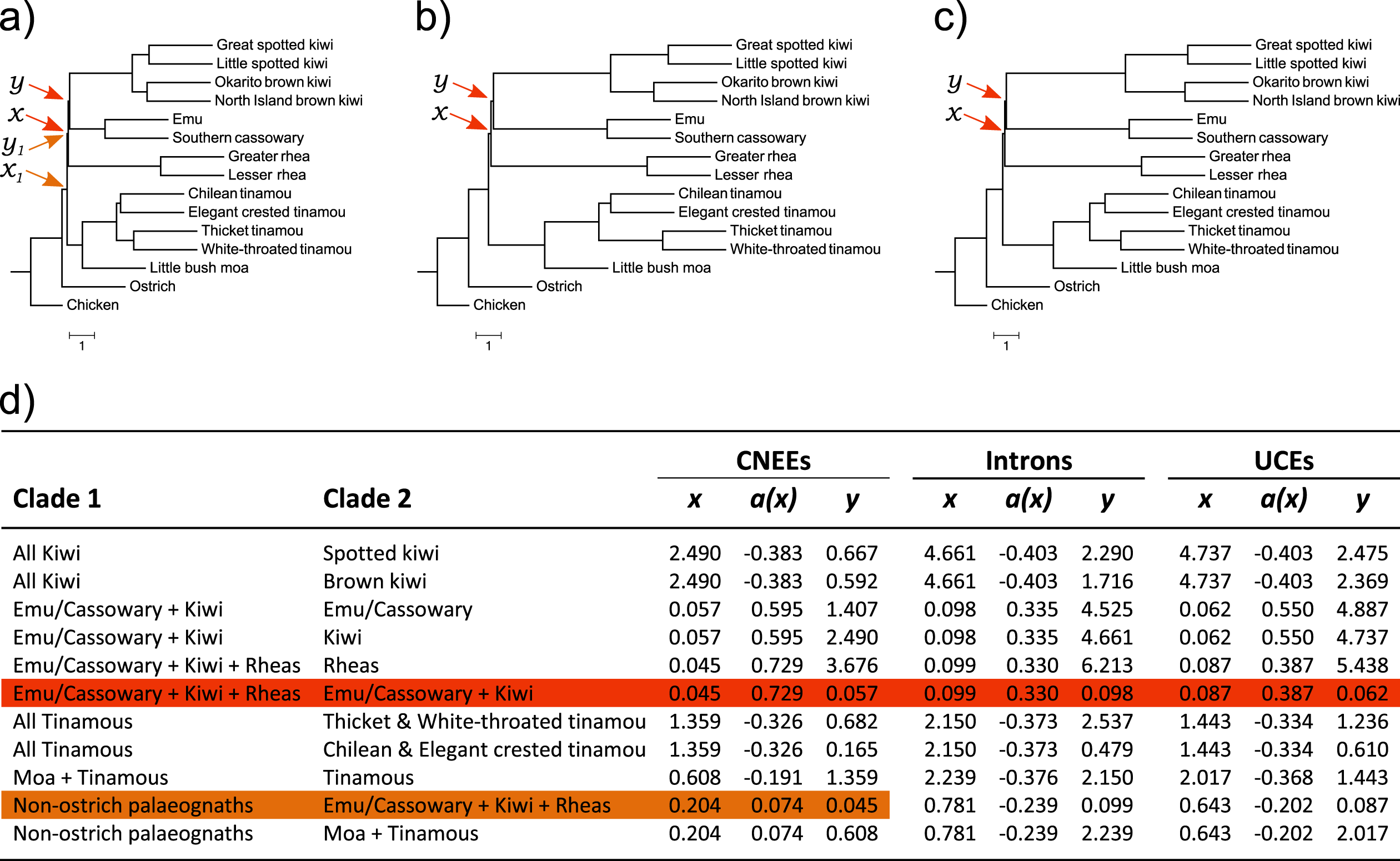
Pairs of short successive internal branches are consistent with expectations for the anomaly zone. Species tree topologies for CNEEs (a), introns (b), and UCEs (c) are shown with internal branch lengths in coalescent units estimated from best maximum likelihood gene trees with MP-EST. Terminal branch lengths are uninformative and are drawn as a constant value across taxa. Coalescent branch lengths for all pairs of branches (x and y) are given in (d), with a(x) calculated following Equation 4 from Degnan and Rosenberg (2006). Anomalous gene trees are expected when y < a(x). Clades fulfilling this anomaly zone criterion are shaded in (d), with the corresponding branches indicated in a-c.

## Discussion

High-throughput sequencing has revolutionized the scale at which phylogenetic inference is being made, facilitating a phylogenomics era with unparalleled opportunity to not only resolve difficult taxonomic relationships but also to better understand the underlying evolutionary processes that have given rise to current biodiversity. In birds, recent large-scale efforts using sequence capture methodologies (McCormack et al. 2013, Prum et al. 2015) and whole-genome sequencing (Jarvis et al. 2014) have helped resolve higher-level interrelationships within Neoaves. Still, some relationships remain poorly resolved or have discordant results across analyses (Suh et al. 2015, Suh 2016, Reddy et al. 2017). Retroposon insertion patterns suggest that a large amount of incomplete lineage sorting (ILS) accompanied a rapid Neoavian radiation (Suh et al. 2015), prompting some authors to propose a hard polytomy involving nine lineages at the base of this group (Suh 2016). Others have stressed the importance of data-type effects in explaining the observed topological incongruence for some clades, and urge researchers to incorporate evidence from as many different marker types as possible when addressing difficult phylogenetic questions (Edwards et al. 2017, Reddy et al. 2017). Together, these results illustrate the advances made with genome-scale data, but also highlight that large amounts of data alone are not necessarily sufficient to resolve deep divergences in the Avian Tree of Life (Philippe et al. 2011).

In parallel, there has been a renewed focus on palaeognath phylogeny throughout the past decade largely motivated by reports that the volant (flighted) tinamous are nested within palaeognaths rather than forming the sister to a clade of flightless ratites (Hackett et al. 2008, Harshman et al. 2008, Phillips et al. 2010, Haddrath and Baker 2012, Baker et al. 2014) and more recent findings that the extinct elephant birds of Madagascar are sister to the New Zealand kiwi (Mitchell et al. 2014, Grealy et al. 2017, Yonezawa et al. 2017). These relationships are consistently recovered across recent molecular phylogenies, but conflicting results for other relationships suggest that ILS across short internal branches in palaeognaths could mirror the discord arising from rapid ancient divergences seen in Neoaves.

We recover a topology placing rheas as sister to emu/cassowary + kiwi that is congruent across all analyses for MP-EST and ASTRAL species tree methods, but that differs from the placement of rheas as sister to the remaining non-ostrich palaeognaths when concatenated data are analyzed. These conflicting topologies are recovered with maximal support in at least some data sets for both coalescent and concatenation analyses. However, subsampling approaches that incorporate resampling across genes as well as sites (double bootstrapping, Seo et al. 2008, Edwards 2016b) provide more robust support for the coalescent species tree, and this topology is further corroborated by patterns of CR1 retroelement insertions from multiway genome-wide screening. However, conflicting CR1 insertions suggest extensive ILS across short internal branches separating major palaeognath lineages, and coalescent lengths for these pairs of branches fall within the theoretical range expected to produce anomalous gene trees. We indeed find that the most common gene tree for each marker type does not match the inferred species tree topology, consistent with an empirical anomaly zone in palaeognaths.

Although we contrast results from concatenation and coalescent species tree methods, we reiterate previous statements that concatenation should be viewed as a specialized case of the multispecies coalescent rather than in strict opposition to it (Liu et al. 2015a,c; Edwards 2016a; Edwards et al. 2016). Concatenation is expected to perform well when coalescent variation among gene trees is low (Liu et al. 2015a,b; Tonini et al. 2015; Mirarab et al. 2016), and we find that most palaeognath relationships are robustly supported by both concatenation and coalescent methods as well as by CR1 insertions. Of particular note, we recover congruent topologies across all sequence-based analyses that nest the tinamous as the sister group to moa within a paraphyletic ratite clade. This moa-tinamou clade is supported even at the level of individual loci, where 94% of both introns and UCEs recover this relationship. Ratite paraphyly is further corroborated by 18 retroelement insertions that are shared by tinamous and non-ostrich palaeognaths, whereas no CR1s supporting ratite monophyly were found.

Unlike the case described above, gene tree heterogeneity resulting from coalescent processes can yield topologies with high support for erroneous clades under concatenation, since this method assumes that all genes evolve with a common topology (Kubatko and Degnan 2007; Liu et al. 2015a,c; Roch and Steel 2015; Edwards et al. 2016). In contrast, the multispecies coalescent models gene trees as conditionally independent variables and, by accommodating gene tree heterogeneity, can accurately infer species tree topologies under high levels of ILS and even in the extreme case of the anomaly zone (Liu et al. 2010, 2015c; Edwards et al. 2016; Mirarab et al. 2016). We believe this fundamental difference in approaches to species tree inference underlies the discordant placements of rheas in our concatenation and coalescent analyses. While recognizing caveats pertaining to possible data-type effects of coding sequence (Reddy et al. 2017), we also note that ASTRAL analysis of anchored hybrid enrichment loci by Prum et al. (2015) produced an identical topology to our coalescent species tree for palaeognaths, whereas Bayesian estimation from concatenated loci was identical to our concatenated analyses of CNEEs and UCEs under maximum likelihood.

Symmetrical counts for the two minor triplet topologies of rheas, emu + cassowary, and kiwi, as well as the symmetry of conflicting retroelement insertions for these alternative hypotheses, suggest a role for ILS in producing gene tree heterogeneity. Importantly, though, both the major triplet topology and statistical tests of CR1 insertions support rheas as sister to emu/cassowary + kiwi within an overarching bifurcating species tree topology as recovered by MP-EST and ASTRAL (Pamilo and Nei 1988, Kuritzin et al. 2016). Retroelement support for the alternative placement of rheas as sister to all other non-ostrich palaeognaths as found by concatenation is entirely absent, whereas CR1 support for a sister-group relationship of emu and cassowary with tinamous, as found under concatenation for CNEEs and UCEs, is strongly rejected in favor of emu/cassowary + kiwi as occurs in all other analyses, including concatenated introns. Instances of retroposon homoplasy through exact parallel insertions or precise deletion are known, but are believed to be rare (Han et al. 2011, Ray et al. 2006, Suh et al. 2015). These results thus strongly corroborate both the inferred species tree topology and the existence of substantial ILS across short internodes separating several palaeognath lineages. We emphasize that the large amount of gene tree heterogeneity in this study is mostly generated by only two short internodes. Our study thus underscores how small numbers of short branches in the species tree can nonetheless generate substantial heterogeneity of largely similar gene trees differing by a small number of nodes.

A question raised for empirical studies is the degree to which gene tree estimation error impacts the performance of summary species tree methods such as MP-EST and ASTRAL that use estimated gene trees as input, since these methods assume gene tree heterogeneity arises from coalescent variation rather than from analytical errors associated with low information content of individual loci, alignment uncertainty, mutational bias, or inappropriate models of sequence evolution (Huang and Knowles 2009, Roch and Warnow 2015, Xi et al. 2015, Blom et al. 2017). We find high average bootstrap support for estimated gene trees and high gene support frequencies for most palaeognath relationships, indicating that marker choice is appropriate to address these questions and gene tree uncertainty is largely confined to the short internal branches inferred to lie within the anomaly zone. Despite this uncertainty, we find that most loci (80-90%) contain variation that is sufficient to support their inferred gene tree topology relative to that of the species tree in likelihood comparisons, and simulations indicate that, for most comparisons, more than 70% of the observed gene tree heterogeneity can be accounted for by coalescent processes alone. While gene tree estimation error certainly occurs in our data set, the consistency of rooted triplet proportions across the three marker types further suggests that, on the whole, estimated gene trees for these data sets provide an adequate and unbiased representation of the coalescent variation in gene tree histories.

Simulations have shown that gene tree summary methods can accurately infer the species tree topology despite gene tree estimation error if the sampling of gene trees is sufficiently large and gene tree inference is unbiased (Liu et al. 2015a,c; Xi et al. 2015; Mirarab et al. 2016; Xu and Yang 2016). Adding weak genes to data sets decreased the performance of summary methods such as MP-EST relative to data sets of strong genes alone, but these methods are nevertheless expected to converge upon the correct estimate of the species tree with an increasing number of genes, even if those genes are minimally informative (Xi et al. 2015, Xu and Yang 2016). These inferences could explain observed patterns for CNEEs, which are on average much shorter and less variable than introns or UCEs and have lower bootstrap support for clades in anomalous gene trees that conflict with the species tree, but which still yield congruent estimates of the species tree topology for coalescent methods. Despite lower per-locus information content, the higher alignment certainty and lower estimated homoplasy of CNEEs relative to other marker types (Edwards et al. 2017) could allow CNEEs to converge upon the correct species tree topology for palaeognaths with high support even for intermediate numbers of loci, but only when methods that accommodate gene tree heterogeneity are used. Simulations investigating the effects of gene tree estimation error also assume there is no bias in gene tree estimation, which can cause summary methods to produce inconsistent results (Xi et al. 2015). We doubt that biased gene trees explain the results for palaeognaths since that explanation would necessitate a similar bias across data sets of three different marker types, would require bias prevalent enough to yield consistent results from even small subsampling replicates within marker types, and would have to generate results that are also consistent with patterns of CR1 insertions.

An important and related question is whether we can confidently detect an empirical anomaly zone given that the same short internal branches expected to produce anomalous gene trees also increase the likelihood of gene tree estimation error due to the short time interval for informative substitutions to accumulate in individual loci of finite length (Huang and Knowles 2009). Using highly variable markers might not ameliorate the situation for deep divergences because of the increased probability for subsequent mutations along long terminal branches to obscure informative substitutions on short antecedent branches (Philippe et al. 2011, Haddrath and Baker 2012, Liu et al. 2015b). Huang and Knowles (2009) demonstrated through simulation that the range of species tree branch lengths producing AGTs expands when the mutational variance of estimated gene trees is considered and that unresolved gene trees, rather than AGTs, would predominate within this region of tree space. This expanded anomaly zone might be what is observed for CNEEs in palaeognaths; although ILS between tinamous and other non-ostrich lineages is clearly evident from CR1 retroelements, only CNEEs infer an anomaly zone in this region of the species tree. In contrast, all three marker types support an anomaly zone across branches separating rheas from emu + cassowary and kiwi. For introns and UCEs, the relative likelihood of anomalous gene trees in most cases is substantially greater than if alignments are constrained to the species tree topology and clades that conflict with the species tree tend to have at least 50% median bootstrap support, suggesting that gene tree heterogeneity is real rather than reflecting gene tree estimation error alone. However, we concur that short internodes pose substantial challenges to accurate gene tree inference, and investigations of the empirical anomaly zone should greatly benefit from algorithms that make ‘single-step’ coestimation of gene trees and species trees scalable to phylogenomic data sets.

In conclusion, we find strong evidence that past difficulty in resolving some palaeognath relationships is likely attributable to extensive incomplete lineage sorting within this group, and that species tree methods accommodating gene tree heterogeneity produce robustly supported topologies despite what appears to be an empirical anomaly zone. We echo the sentiments of other authors that high bootstrap support alone is an inadequate measure of confidence in inferred species trees given increasingly large phylogenomic data sets (Edwards 2016b, Sayyari and Mirarab 2016). Congruence across marker types and corroboration from rare genomic changes such as retroposon insertions, as well as phylogenomic subsampling strategies to assess the underlying phylogenetic signal, will increase our confidence in recovered topologies and could also highlight which conflicts are robustly supported and merit further investigation (Tonini et al. 2015, Edwards 2016b, Mirarab et al. 2016, Reddy et al. 2017). Congruence is certainly not a new idea in phylogenetics, but an increasing emphasis on its importance in the era of species trees will continue to advance the field beyond reports of highly supported, but often discordant, ‘total evidence’ topologies toward a more nuanced ‘sum of evidence’ approach that considers not just which topologies are produced but also what they can tell us about the underlying evolutionary processes and our attempts to model them.

## Acknowledgement

We thank Liang Liu for advice regarding coalescent simulation, and members of the Edwards lab for helpful discussions. All analyses were run on the Odyssey cluster supported by the FAS Division of Science, Research Computing Group at Harvard University. Support was provided by NSF grant DEB 1355343 (EAR 1355292) to AJB, MC, and SVE.

---

**Suppl. Figure S1** Palaeognath phylogenetic relationships inferred with MP-EST (a-d) and ASTRAL (e-h) coalescent-based species tree methods and maximum-likelihood inference of fully partitioned concatenated alignments with ExaML (i-l). Topologies were inferred from 12,676 conserved non-exonic elements (CNEEs), 5,016 introns, 3,158 ultraconserved elements (UCEs), and the total evidence data set of 20,850 loci combined across marker types (TENT). Bootstrap supports are drawn for clades with < 100% support.

**Suppl. Figure S2** Support for alternative hypotheses of the sister group to emu + cassowary from phylogenomic subsampling using MP-EST (a-c), ASTRAL (d-f), and ExaML (g-i). Plots display the mean bootstrap support for each hypothesis from 10 replicates of randomly sampled loci within each data set size category (e.g. 50-3000 loci, shown on x-axis).

**Suppl. Figure S3** Heatmap of the number of replicates from phylogenomic subsampling that support each alternative hypothesis for the sister group to emu + cassowary at a minimum bootstrap support of 70% (a-i) and 90% (j-r). Rows labelled H1-H4 within each panel correspond to the four alternative hypotheses outlined in the legend. Columns within each panel labelled 50-3000 indicate the number of loci subsampled at random with replacement from all loci for each marker type. Coloring of cells indicates the number of replicates (of 10 in total for each data set size category) that support each hypothesis at the given bootstrap cut off, corresponding to the colouring scheme outlined in the legend.

**Suppl. Figure S4** Support for the four most common gene tree topologies for each marker type. Median bootstrap support (a-c) and the median number of substitutions under a parsimony criterion (d-f) for branches in gene tree topologies that conflict with the species tree. Letters below each group refer to topologies shown in (j), with numbers referring to branches labelled on those topologies. Reference lines in (a-c) are drawn at 50% bootstrap support. (g-i) Violin plots of ΔAIC for loci belonging to each AGT category. Values show the difference in AIC for sequence alignments given the majority rule extended consensus gene tree topology, relative to the AIC when sequence alignments are constrained to the species tree topology. A reference line is drawn at ΔAIC(_gene tree-species tree_)= -2, with values beneath this cut off indicating substantially stronger support in favor of the gene tree topology (Burnham and Anderson 2002).

**Suppl. Figure S5** Summary measures of aligned sequence data for each marker type (CNEEs, introns, UCEs), showing the total aligned sequence length per locus (a-c), variable alignment columns as a percent of the total alignment length (d-f), and the percent of parsimony informative alignment columns per locus (g-i).

**Suppl. Figure S6** Pairs of short successive internal branches are consistent with expectations for the anomaly zone. Species tree topologies for CNEEs (a), introns (b), and UCEs (c) are shown with internal branch lengths in coalescent units estimated from best maximum likelihood gene trees with ASTRAL. Terminal branch lengths are uninformative and are drawn as a constant value across taxa. Coalescent branch lengths for all pairs of branches (x and y) are given in (d), with a(x) calculated following Equation 4 from Degnan and Rosenberg (2006). Anomalous gene trees are expected when y < a(x). Clades fulfilling this anomaly zone criterion are shaded in (d), with the corresponding branches indicated in a-c.

